# Improved 3D Radial Phyllotaxis Trajectories for Uniform Density Distribution of Readout Directions and Sequential Binning

**DOI:** 10.64898/2026.07.06.736720

**Authors:** Mauro Leidi, Jim Délitroz, Eva Peper, Yiwei Jia, Jaime Barranco, Jean-Baptiste Ledoux, Ludovica Romanin, Jessica A.M. Bastiaansen, Juliane Schneider, Benedetta Franceschiello

## Abstract

**Purpose:** To develop 3D radial spiral phyllotaxis trajectories that provide a uniform density distribution of readout directions and support retrospective sequential binning, thereby reducing ringing artifacts and improving image quality.

**Methods:** UPhy trajectory redefines the polar angle to achieve uniform density distribution of readout directions. FlexiPhy further decouples the azimuthal and polar ordering of interleaves through a randomized permutation, improving robustness to sequential binning. The proposed trajectories were evaluated in vivo on 10 healthy volunteers using two gradient-echo sequences on a 3T MRI scanner. Sequential temporal reconstructions were compared with reference reconstructions using structural similarity and relative L2 error metrics.

**Results:** UPhy presents analytically demonstrated uniform density distribution of readout directions. Quantitative analysis shows significantly higher SSIM values and lower relative L2 errors for FlexiPhy compared with both the original phyllotaxis and UPhy trajectories after Bonferroni correction (*p*_corrected_ < 0.05).

**Conclusion:** FlexiPhy enables more reliable sequential binning reconstructions by reducing trajectory-induced ringing artifacts and temporal inconsistencies. Moreover, its randomized construction is not tied to a specific binning strategy, making it broadly compatible with retrospective binning approaches used in dynamic and motion-resolved MRI.

## 1 | INTRODUCTION

Three-dimensional radial trajectories, such as spiral phyllotaxis ^1^, provide motion robustness, self-navigation capabilities, and flexible retrospective data sorting (or binning), making them attractive for dynamic and motion-resolved MRI applications including cardiac imaging ^2,3,4,5,6^, fetal cardiac imaging ^7^, lung imaging ^8,9,10^, liver imaging ^11^, dynamic eye imaging ^12,13,14^, neuroimaging ^15,16^, real-time applications such as speech imaging ^17^, as well as Zero Echo Time ^18^ and Ultra-short Echo Time imaging applications ^19,20,21,22^. Spiral phyllotaxis further combines efficient k-space coverage with smooth transitions between successive readouts, reducing eddy-current effects ^1,23,24,25^while enabling frequent superior–inferior navigator acquisitions for motion tracking and self-navigation ^26^.

For non-Cartesian acquisitions, sampling efficiency is governed by the largest gap between neighboring k-space samples ^25,27^. Although radial trajectories intentionally oversample the center of k-space, where most of the signal energy is concentrated, a uniform spherical distribution of sampling density (SDSD) ^28,21^, or equivalently a uniform density distribution of readout directions, remains desirable. A uniform SDSD minimizes large sampling gaps, thereby reducing sampling inefficiencies, mitigating aliasing artifacts, and improving signal-to-noise ratio (SNR) ^28,29,30,31^. Conversely, non-uniform SDSD produces anisotropic k-space sampling, leading to direction-dependent variations in effective field of view and spatial resolution ^32,33^. Recent studies have shown that the spiral phyllotaxis trajectory does not achieve uniform SDSD ^32,34^. Furthermore, single-hemisphere sampling may increase sensitivity to trajectory-dependent phase inconsistencies, which can be mitigated by including readouts from both hemispheres ^35^.

While global SDSD characterizes the overall trajectory, many applications reconstruct only subsets of the acquired readouts. In these settings, the sampling properties within reconstruction bins become equally important ^21^. Spiral phyllotaxis has been successfully employed in free-running cardiac MRI ^4^, where retrospective bins are formed by selecting readouts acquired throughout the entire scan. As a result, each bin retains a well-distributed set of sampling directions and favorable k-space coverage despite containing only a subset of the acquired data. However, some applications require to bin data into temporally contiguous subsets of readouts, for example when reconstructing dynamic image series, functional MRI applications ^16^, or performing rigid motion estimation for fetal cardiac imaging ^7^. However, consecutive subsets of spiral phyllotaxis acquisitions have been shown to produce clustered sampling patterns and non-uniform k-space coverage ^36^. These sampling irregularities have been associated with ringing artifacts in reconstructed images ^28^, potentially degrading image quality and affecting downstream processing tasks that rely on temporally resolved reconstructions, including motion estimation and correction.

Consequently, an ideal phyllotaxis-based trajectory should provide both uniform SDSD and robustness to arbitrary retrospective binning strategies. Spiral phyllotaxis achieves efficient global k-space coverage and favorable gradient behavior but do not explicitly address either the non-uniform SDSD or the deterioration of sampling characteristics within temporally contiguous reconstruction bins.

This work introduces two methodological advancements that progressively improve the sampling characteristics of the phyllotaxis trajectory. First, Uniform Phyllotaxis (UPhy) modifies the polar-angle formulation of the original trajectory to improve density distribution of the readout directions. Building upon this formulation, Flexible Phyllotaxis (FlexiPhy) introduces a randomized ordering strategy that decouples the polar and azimuthal organization of interleaves, thereby improving the distribution of readouts within sequential reconstruction bins while preserving the favorable properties of the original design. The purpose of this work was to evaluate the sampling characteristics and image reconstruction performance of UPhy and FlexiPhy relative to conventional spiral phyllotaxis.

## 2 | METHODS

### 2.1 | Standard phyllotaxis model

In radial trajectories, a readout is typically a straight line in k-space passing through the center, along which equidistant points are sequentially sampled, each of which is associated to a measurement in the form of a complex value in the frequency domain. The first sample of each readout, called its origin, lies on the surface of a half-sphere (or, in some cases, a full sphere), as illustrated in the left panel of Figure 1. The collection of such origins is referred to here as the sampling pattern, which determines the SDSD of the trajectory. One can analyze the geometrical properties of the sampling pattern, which can then be extended radially to all measurements by exploiting symmetries. The final trajectory specifies also the order in which the readouts themselves are acquired.

**FIGURE 1.**
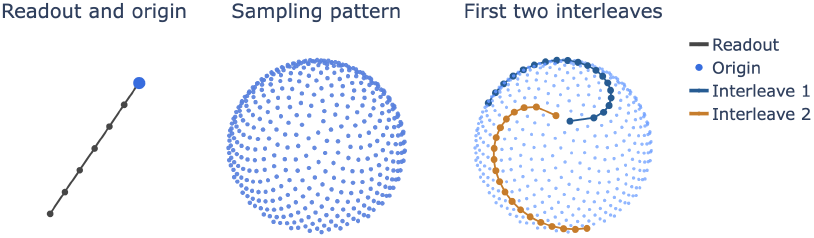
Illustration of the spiral phyllotaxis trajectory. Left: single radial readout line and its origin. Center: the distribution of the points in the sampling pattern over the half-sphere ensures no overlapping. Right: the first two interleaves are highlighted, illustrating the golden-angle interleaving strategy and the small angular distance between successive readouts within an interleave.

#### 2.1.1 | Definition of the Spiral Phyllotaxis

The spiral phyllotaxis sampling pattern is the three-dimensional generalization of Vogel’s model of a planar spiral ^37^. For a sampling pattern on *N* points, the polar coordinates of point *y*_*n*_, 1 ≤ *n* ≤ *N*, are given by:

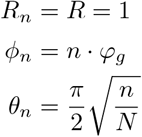

where

- The polar angle *θ*_*n*_ is the angle from the positive *z*-axis of readout number *n*.
- The azimuthal angle *ϕ*_*n*_ is the angle on the *xy*-plane from the positive *x*-axis in the clockwise direction of readout number *n*.
- The radius *R* is the (here constant) distance to the origin.
- The golden angle *φ*_*g*_ is defined as 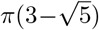 radians.

#### 2.1.2 | Interleaving

Interleaving is a convenient way to order the readouts of the sampling pattern. Readouts are grouped in subsets called interleaves, separated by the golden angle, for two main purposes. First, unlike the main definition of the sampling pattern, it ensures that the trajectory covers different angular directions efficiently, thereby improving the uniformity of readout directions coverage over small time intervals. Second, by grouping readouts with small angular differences, it limits rapid gradient changes between successive readouts, which helps reduce eddy current artifacts. This property is linked to the golden angle *φ*_*g*_. It has been observed that for a Fibonacci number *F*, then *F* · *φ*_*g*_ ≈ 0 mod 2*π* ^38^. As a consequence, the readouts *y*_*n*_ and *y*_*n*+*F*_ will be very close to each other with respect to the azimuthal angle. It is therefore natural to regroup the *N* points of the spiral in *F* interleaves of size *P*, where *N* = *F* · *P* and *F* belongs to the Fibonacci sequence. Interleaves are built so that the *k*-th interleaf comprises the points indexed by {*k, k* + *F, k* + 2*F*, …, *k* + (*P* − 1)*F*}. Figure 1 illustrates the complete sampling pattern together with the first two interleaves, highlighting both the efficient angular coverage induced by the golden-angle ordering and the small angular distance between successive readouts within each interleave. However, in practice it’s recommended not to select *F* as a large Fibonacci number, but to select a combination of *F* and *P* such that 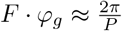 mod 2*π*.

The explicit polar coordinates of 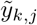 the *j*-th point in interleave *k* according to the standard phyllotaxis scheme (equivalently, the (*k* +(*j* − 1)*F*)-th point of the sampling pattern) are:

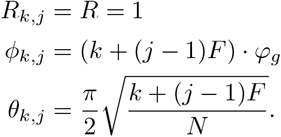

During acquisition, readouts are ordered by interleave, so that the points of interleave 1 are acquired first, followed by interleave 2, and so on.

### 2.2 | UPhy: an acquisition scheme with uniform SDSD

In the standard phyllotaxis model, the polar angle does not yield a perfectly uniform distribution of points over the surface of the half-sphere, but instead tends to cluster more points near the equator as presented in Figure 2. A simple redefinition of the polar angle to:

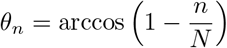

defines UPhy, a trajectory that is very similar to the standard phyllotaxis. The azimuthal angles are unchanged, while the polar angles still span the range from 0 to 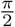, but the point distribution becomes uniform over the sphere, preventing clustering along the equator, as shown in Figure 2. Quantitative explanations and an analytical proof of uniform SDSD are provided in section 8 of the supplementary material.

**FIGURE 2.**
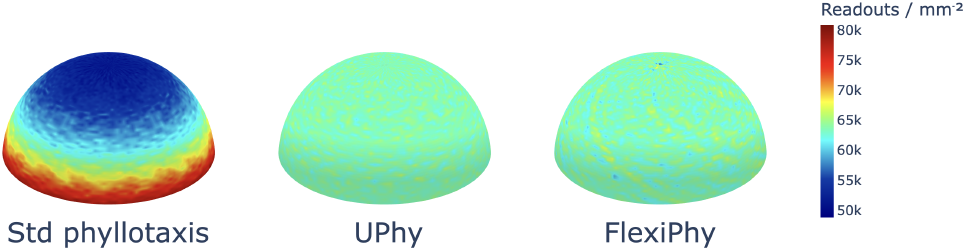
Sampling density of readouts for the three trajectories on the half-sphere. **(Left)** The standard phyllotaxis trajectory exhibits a clear density inhomogeneity, with increased sampling concentration near the equatorial region. **(Center)** UPhy substantially improves the uniformity of the readout direction distribution density by correcting the polar-angle bias. **(Right)** The SDSD of FlexiPhy is similar to UPhy’s. Densities are displayed using a shared global color scale, allowing direct quantitative comparison between trajectories.

### 2.3 | FlexiPhy : an acquisition scheme for flexible binning

In the case of the standard phyllotaxis, groups composed of only *f* < *F* (for example, *f* = *F/*4) consecutive inter-leaves induce significantly non-uniform SDSD in each bin. More specifically, when looking at the readout origins, circular gaps can be observed in the k-space, which arise as a consequence of the phyllotaxis definition, as illustrated in Figure 3. After reconstruction, those gaps manifest as ringing artifacts, as described by Ding & al.^28^.

**FIGURE 3.**
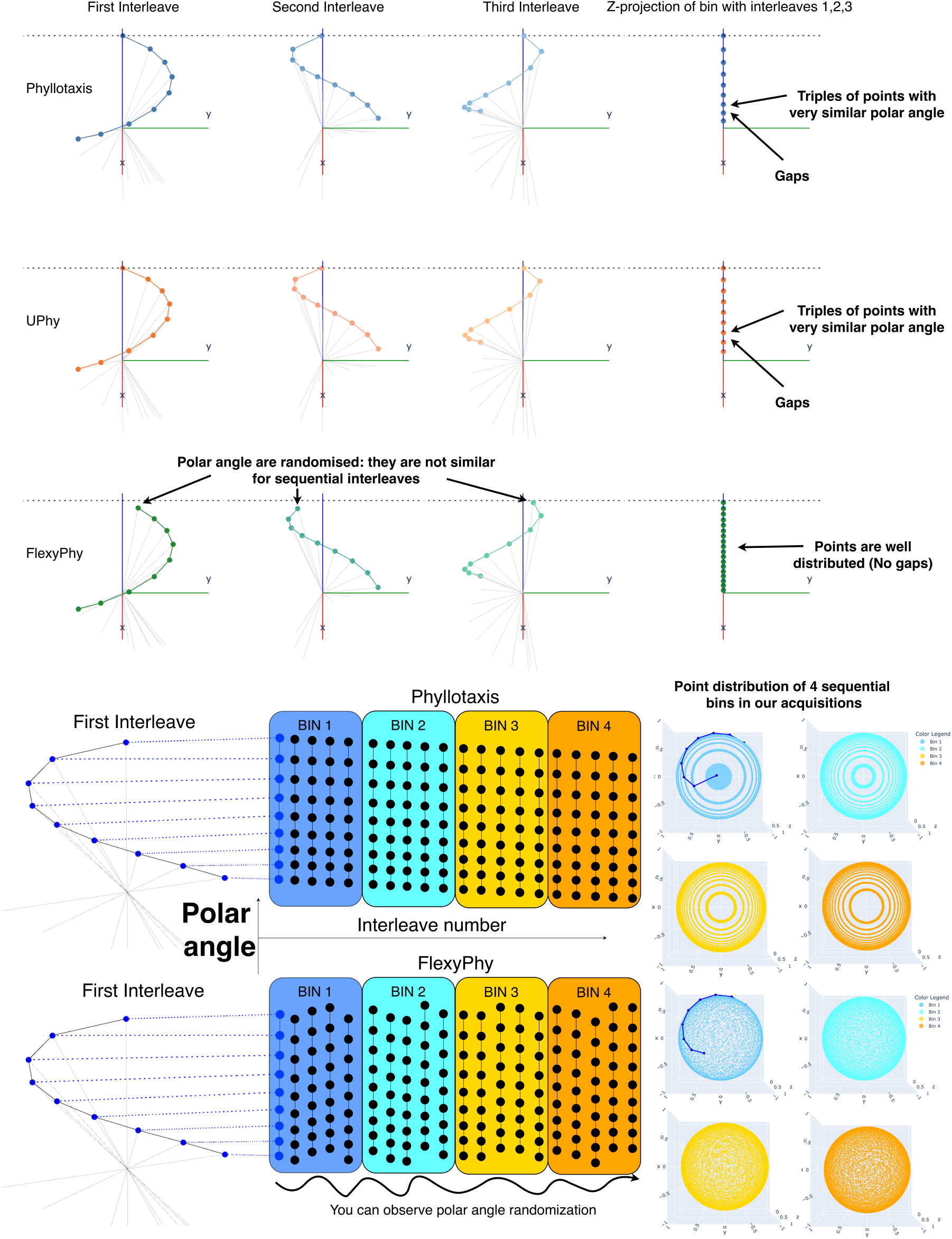
The standard phyllotaxis and UPhy spirals both show a monotonically decreasing polar coordinate with increasing interleaf number, resulting in a point accumulation that appear as concentric rings when using sequential binning. FlexiPhy mitigates this effect through polar angle randomization, resulting in a drastically improved point distribution when sequential binning is used.

UPhy can be further improved by decoupling the azimuthal and polar angles. This preserves the golden-angle separation between successive interleaves while avoiding the systematic clustering along the *z*-axis that leads to ringing artifacts in sequential binning (Figure 3). The resulting trajectory is defined by the following spherical coordinates of the *j*-th point in the *k*-th interleaf. Given a permutation *σ* achieving the decoupling, and using the polar angle definition introduced in subsection 2.2:

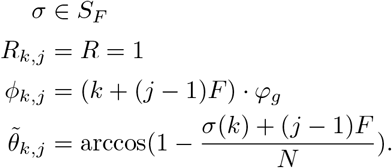

where *S*_*F*_ is the symmetric group on *F* elements, so that *σ* is a randomly selected permutation of our *F* interleaves, see section 8 of the supplementary material for precautions on how to use this formulation and the statistical properties linked to this randomization.

### 2.4 | Data Acquisition

Ten healthy volunteers underwent a single MRI session on a 3T clinical scanner (MAGNETOM Prisma Fit, Siemens Healthcare, Forchheim, Germany) equipped with a 64-channel head/neck coil. This study received approval from the CER-VD cantonal ethics committee (Project-ID 2023 − 02197). All volunteers provided written informed consent prior to participation and for the release of their data in a public data repository. Visual stimuli were back-projected onto a mirror attached to the head coil, with a total distance between the participants’ eyes and the screen of 102 cm. The visual field on the mirror spanned 18°.

In the scanner, every participant was asked to accomplish two distinct tasks. During the first task, the participant had to stare at a central fixation stimulus, composed of two circles and a cross. This fixation shape is designed to reduce undesired eye movements, and was selected based on the results reported by Thaler et al. ^39^. This stimulus was presented to make sure the participant keeps the eyes as still as possible. The second task, is a simplified version of the protocol presented in ^12^, which consists in dynamic imaging of the moving eye. The goal of the second task is to present a second acquisition in a more realistic dynamic imaging scenario and assess the robustness across imaging sequences. In this setup, the fixation shape was presented at four equidistant sequential locations along the horizontal direction, from left to right. Both stimuli were implemented using Psychopy ^40^ and stimulation code is available on GitHub. The same program enables the calibration of the eye-tracker from outside the scanner room. Each participant repeated both tasks three times, for the data to be acquired using different trajectories (the standard phyllotaxis, Uphy and FlexiPhy). An eye-tracking (ET) system (EyeLink 1000Plus, SR Research) was used to record eye movement trajectories (of the right eye) using infrared light, with a sampling rate of 1000 Hz, through an infrared compatible mirror positioned inside the scanner bore. A NordicLabs SyncBox was used to synchronize the scanner, the ET, and the visual stimulation. A complete description of the hardware setup and acquisition can be found in the standard operating procedures linked to this study.

Data were acquired using two uninterrupted 3D gradient-recalled echo (GRE) sequences, the first consisting of a T1-weighted and the second corresponding to a fat-suppressed T1-weighted using lipid-insensitive binomial off-resonant RF excitation pulses ^41^ (LIBRE). Each sequence was combined with the three different 3D radial sampling trajectories (the standard phyllotaxis, Uphy and FlexiPhy) and was implemented using the Pulseq framework ^42^. The code used to generate the corresponding .seq files is open-source and available on GitHub. The first sequence, i.e. the conventional T1-weighted GRE acquisition, was acquired with the following parameters: repetition time (TR) = 4.5 ms, echo time (TE) = 2.0 ms, flip angle = 12°, bandwidth = 496 Hz/pixel, and isotropic spatial resolution of 1 mm over a 120 × 120 × 120 mm^3^ field of view. Each readout consisted of 240 samples due to a radial oversampling factor of 2. Quadratic RF spoiling was applied with a phase increment of 50°, and the gradient spoiling area was matched to the readout gradient area, a setting determined during the pilot phase to optimize the trade-off between acquisition time and image quality. The second sequence was a modified GRE acquisition employing LIBRE RF excitation pulses to improve visualization of the orbital anatomy, in particular the optic nerve, which is surrounded by adipose tissue. Compared to the conventional T1-weighted GRE sequence, this acquisition used a longer TR and TE (TR = 6.20 ms, TE = 3.62 ms) and the flip angle was reduced to 6°, maintaining an effective flip angle of 12°. All other imaging parameters were matched to the T1-weighted GRE acquisition. Table 1 contains a summary of all acquisition parameters. For both sequences, acquisitions were performed using three different radial trajectory designs, and a superior-inferior navigator was acquired for each interleave. Each dataset consisted of a total of N=82960 radial readouts, organized into F=2440 inter-leaves of P=34 readouts each. In addition to the main acquisitions, for both sequences, two calibration prescans were acquired to enable coil sensitivity estimation. These prescans consisted of N=9218 radial readouts (F=233, P=34) and were acquired using the same trajectory design as the main acquisition. One calibration prescan was performed using the body coil as the receiver, while the second used the receiver 64-channel head/neck coil employed for the data acquisition.

**TABLE 1.** Acquisition parameters for the T1-weighted GRE and LIBRE sequences.

| Parameter | T1-weighted GRE | LIBRE |
| --- | --- | --- |
| Sequence type | 3D GRE | 3D GRE with LIBRE RF pulse |
| TR / TE | 4.5 / 2.0 ms | 6.20 / 3.62 ms |
| Flip angle | 12° | 6° |
| Effective flip angle | 12° | 12° |
| Bandwidth | 496 Hz/pixel | 496 Hz/pixel |
| Spatial resolution | 1 mm isotropic | 1 mm isotropic |
| Field of view | 120 × 120 × 120 mm<sup>3</sup> | 120 × 120 × 120 mm<sup>3</sup> |
| Samples per readout | 240 | 240 |
| Radial oversampling factor | 2 | 2 |
| RF spoiling | Quadratic, 50° increment | Quadratic, 50° increment |
| Spoiling gradient | Matched to readout gradient area | Matched to readout gradient area |
| **Parameters common to both sequences** |  |  |
| Scanner | 3T MAGNETOM Prisma Fit |  |
| Coil | 64-channel head/neck coil |  |
| Trajectories | Standard phyllotaxis, UPhy, FlexiPhy |  |
| Readouts per dataset | $N = 82960$ | |
| Interleaves | $F = 2440$ | |
| Readouts per interleaf | $P = 34$ | |
| Navigator | One superior-inferior readout per interleaf |  |
| Calibration prescans | $N = 9218$ readouts, $F = 233$, $P = 34$ | |

### 2.5 | Image Reconstruction and Analysis

Before image reconstruction, the raw data and the trajectory were corrected for rigid motion. To estimate the motion parameters, a series of low-resolution images was first reconstructed with 2 mm isotropic spatial resolution and 3.5 s temporal resolution. Because these images were used solely for motion estimation, a compressed-sensing reconstruction enforcing similarity between neighboring temporal frames was employed to improve image quality at the high temporal resolution. Rigid motion parameters were then estimated from these images using SPM ^43^. The resulting motion parameters were linearly interpolated in time to assign one motion estimate to each readout. These interpolated estimates were used to correct the trajectory by applying the inverse of the estimated rotation, while k-space measurements were corrected using the shift theoremu ^44^. The motion-corrected data were subsequently reconstructed with a zero-padded gridded reconstruction. Coil sensitivity maps were estimated from prescan calibration data. All reconstruction were performed using the Monalisa toolbox ^45^.

For each of the three trajectories, all acquired readouts were used to reconstruct a reference image. Because this reconstruction uses the complete sampling pattern, it provides the most uniform k-space coverage and serves as the ground truth for subsequent comparisons. In addition, four sequential temporal images were reconstructed, each using one quarter of the acquired readouts, to emulate a sequential-binning scenario. Each temporal-bin reconstruction contained approximately 20,740 readout lines, exceeding the Nyquist requirement of a corresponding Cartesian acquisition (120 · 120 = 14, 400 lines). This choice ensured that any observed image degradation would primarily arise from non-uniform spatial distribution of the readouts rather than from undersampling. Ideally, each temporal image should closely match the corresponding reference image. Similarity between the magnitude images of the temporal bins and the corresponding reference image was quantified using the Structural Similarity Index Measure (SSIM) and the relative L2 error.

For each subject and trajectory, the metrics were first computed separately for the four bins and subsequently averaged to obtain a single subject-level score per metric and trajectory. Statistical differences between trajectories were assessed using paired t-tests on these subject-level scores for all pairwise trajectory comparisons. To account for multiple comparisons, p-values were corrected using the Bonferroni method. Improvements were defined as lower L2 distance and higher SSIM values.

To evaluate the improvement of SDSD provided by UPhy we computed the two uniformity metrics originally reported by Piccini & al.^1^: the average distance between successive points within a given interleaf and the Relative Standard Deviation (RSD) of the distances between each point and its four nearest neighbors. A detailed description of these measures is provided in subsection 8.4 of the Supplementary Material.

## 3 | RESULTS

Figure 5 compares the standard phyllotaxis and UPhy trajectories using the two uniformity metrics originally reported by Piccini & al.^1^: UPhy either matches or improves upon the standard phyllotaxis trajectory according to both metrics.

**FIGURE 4.**
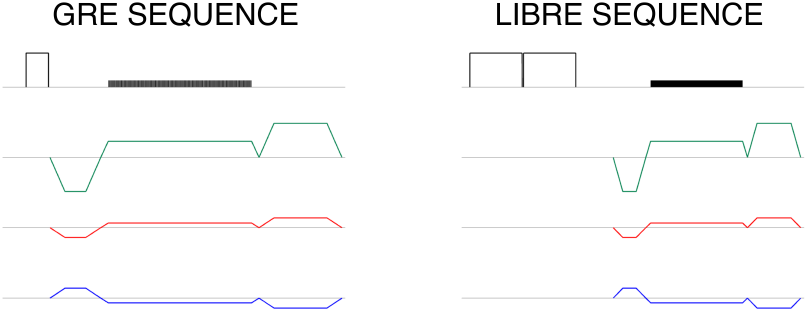
Sequence diagram of a single block for both the T1-weighted GRE and LIBRE sequences. An RF excitation pulse is followed by a prephaser gradient, a readout gradient, and a spoiler gradient. RF spoiling with quadratic increment is also adopted.

**FIGURE 5.**
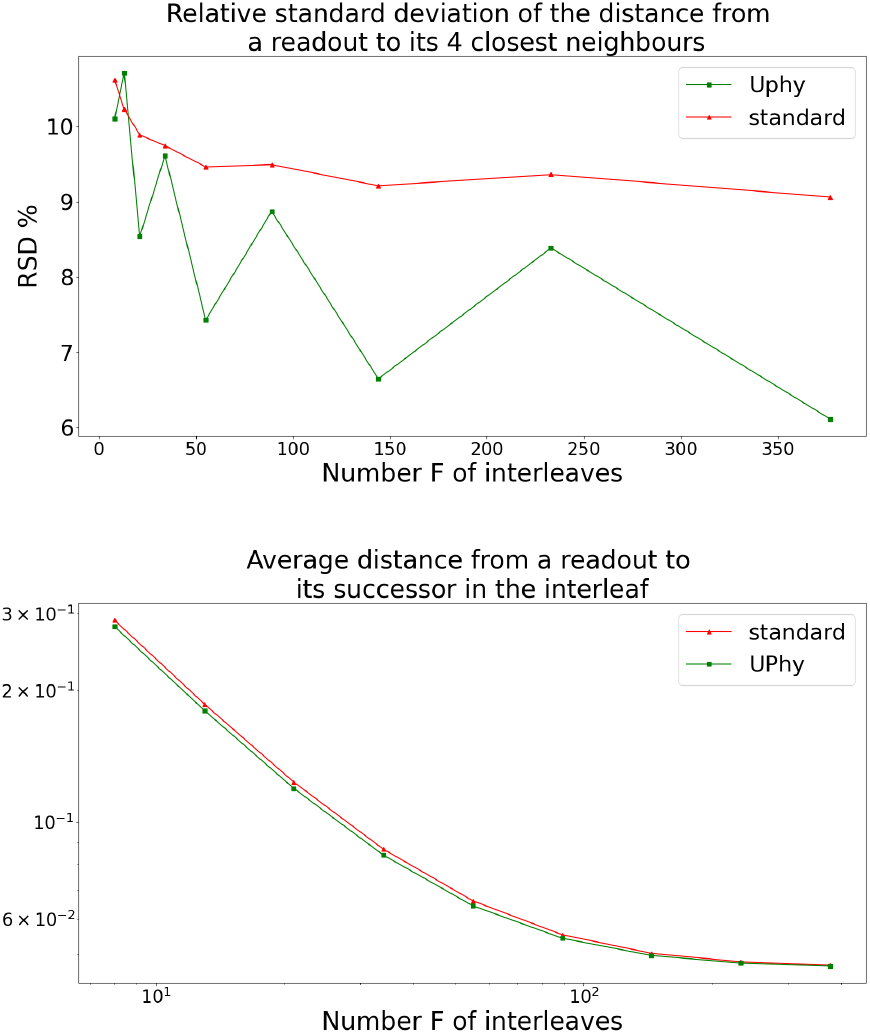
Reproduction of Fig. 3 and 4 of ^1^. According to the two metrics introduced in ^1^, the UPhy trajectory either outperforms or equalizes the original spiral phyllotaxis model.

### 3.1 | Static brain images

An interactive visualization of all reconstructions is publicly available online. Figure 6 shows representative temporal-bin reconstructions and corresponding difference maps relative to the single-bin reference reconstruction for both the GRE and LIBRE acquisitions. For the standard phyllotaxis and UPhy trajectories, sequential-bin reconstructions exhibited visible ringing artifacts, which resulted in temporal inconsistencies across bins. In contrast, the FlexiPhy trajectory substantially reduced these artifacts, producing more temporally stable reconstructions with improved consistency across bins. Difference maps relative to the single-bin reference further highlight these observations.

**FIGURE 6.**
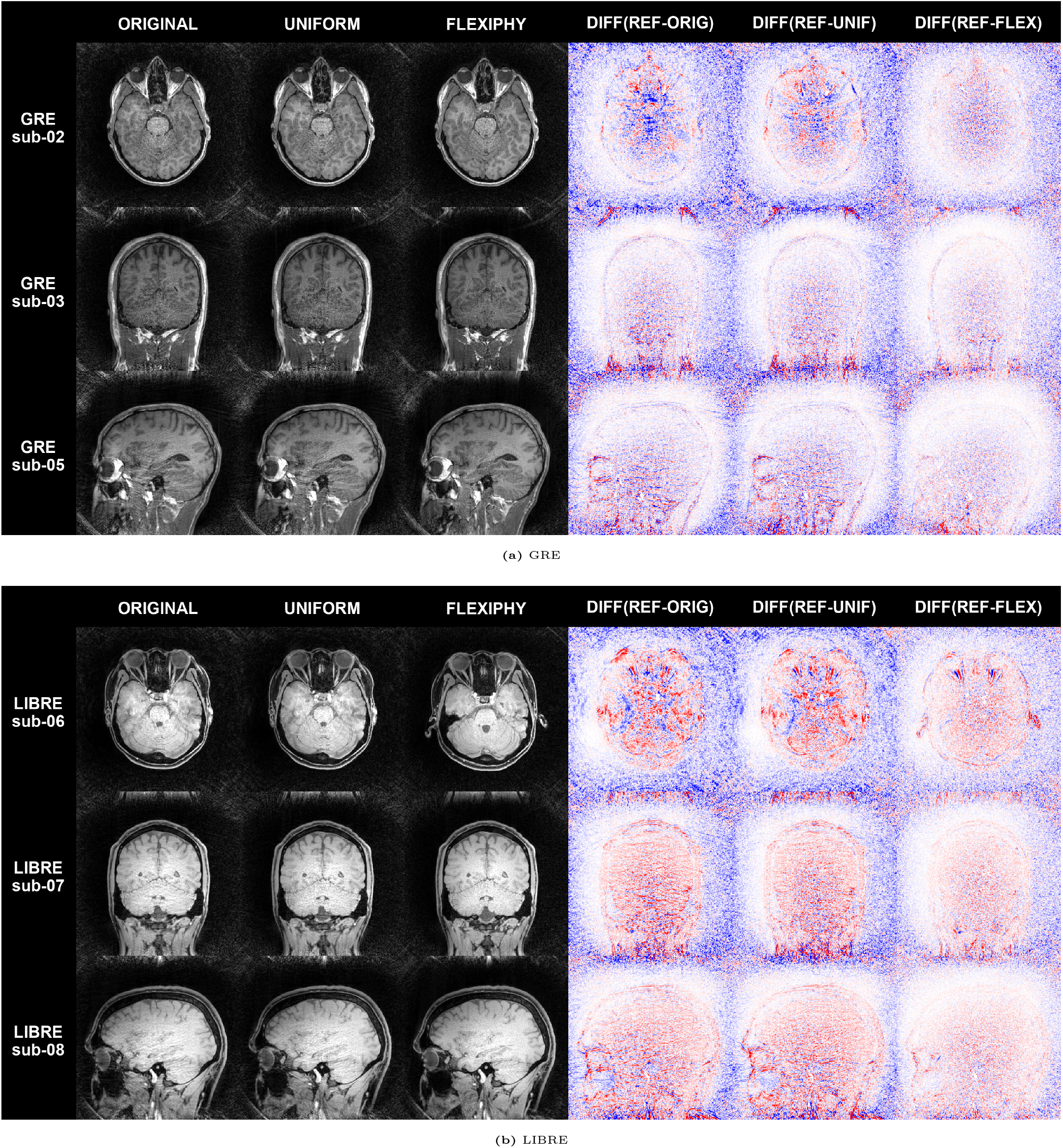
Trajectory-wise comparison of reconstructed temporal bins and corresponding difference maps relative to the single-bin reference reconstructed using all the acquired data.

The quantitative evaluation of the sequential temporal-bin reconstructions shown in Figure 8, based on SSIM and relative L2 distance relative to the corresponding reference images, confirmed the visual observations.

**FIGURE 7.**
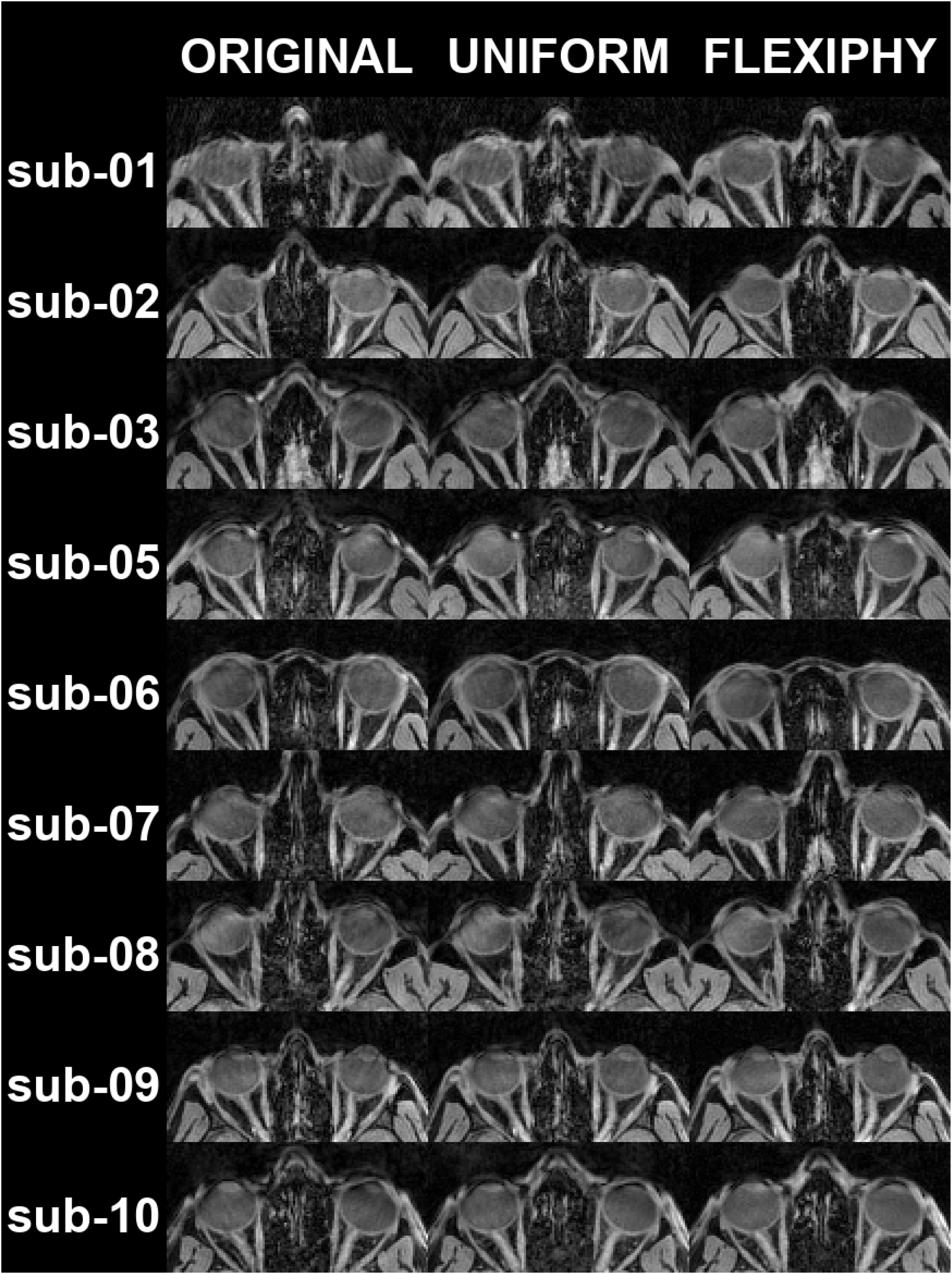
Animated LIBRE axial eye-region collage across the four temporal bins. Similarly to Figure 4, the artifacts due to the trajectory are resolved in the FlexiPhy images.

**FIGURE 8.**
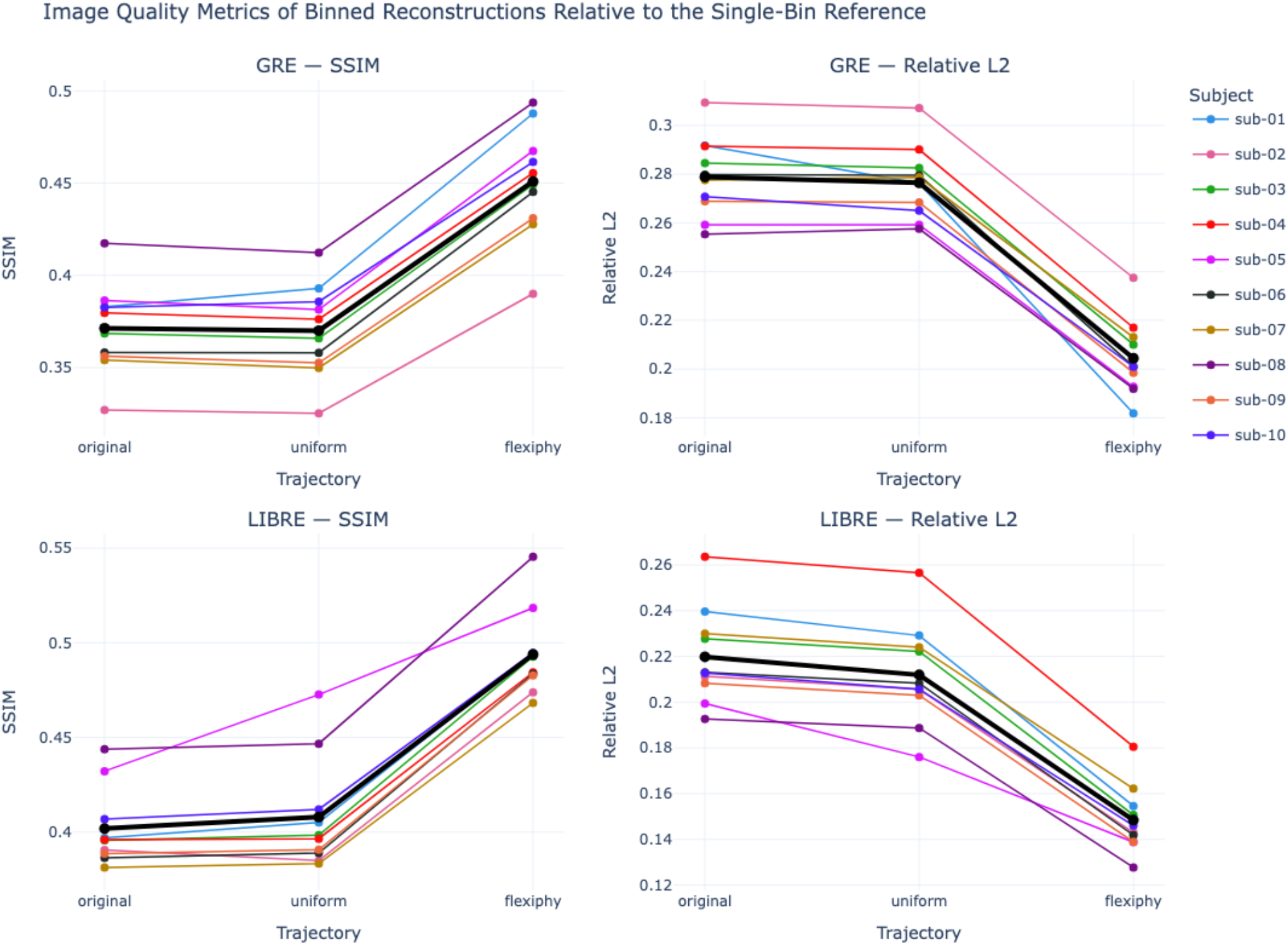
Image-quality metrics for T1-weighted GRE and LIBRE acquisitions across the three acquisition trajectories. For each subject and trajectory, SSIM and relative L2 distance were computed separately for the four bins using the corresponding reference image and then averaged across bins to obtain one subject-level score. Lower relative L2 distance and higher SSIM indicate better agreement with the reference image. Across both sequences, FlexiPhy shows improved agreement with the reference compared with the standard Phyllotaxis and UPhy trajectory.

As summarized in Table 2, pairwise comparisons were performed using paired t-tests on the subject-level scores, and p-values were corrected for multiple comparisons using the Bonferroni method. FlexiPhy yielded significantly higher SSIM values and lower relative L2 distances than both the standard phyllotaxis and UPhy trajectories for both GRE and LIBRE GRE acquisitions. For GRE, FlexiPhy significantly improved SSIM compared with the standard trajectory (*p*_corr_ = 2.90 × 10^−8^) and UPhy (*p*_corr_ = 1.66 × 10^−9^), while also reducing the relative L2 distance compared with the standard trajectory (*p*_corr_ = 3.28×10^−7^) and UPhy (*p*_corr_ = 1.39×10^−8^). Similar results were observed for LIBRE GRE, where FlexiPhy significantly improved SSIM relative to the standard trajectory (*p*_corr_ = 5.38 × 10^−11^) and UPhy (*p*_corr_ = 2.47 × 10^−7^), and significantly reduced the relative L2 distance (*p*_corr_ = 4.60 × 10^−9^ and 2.23 × 10^−7^, respectively). No significant differences in SSIM were observed between the standard phyllotaxis and UPhy trajectories. The only statistically significant difference between these two trajectories was a lower relative L2 distance for UPhy in the LIBRE GRE acquisition (*p*_corr_ = 0.021).

**TABLE 2.**
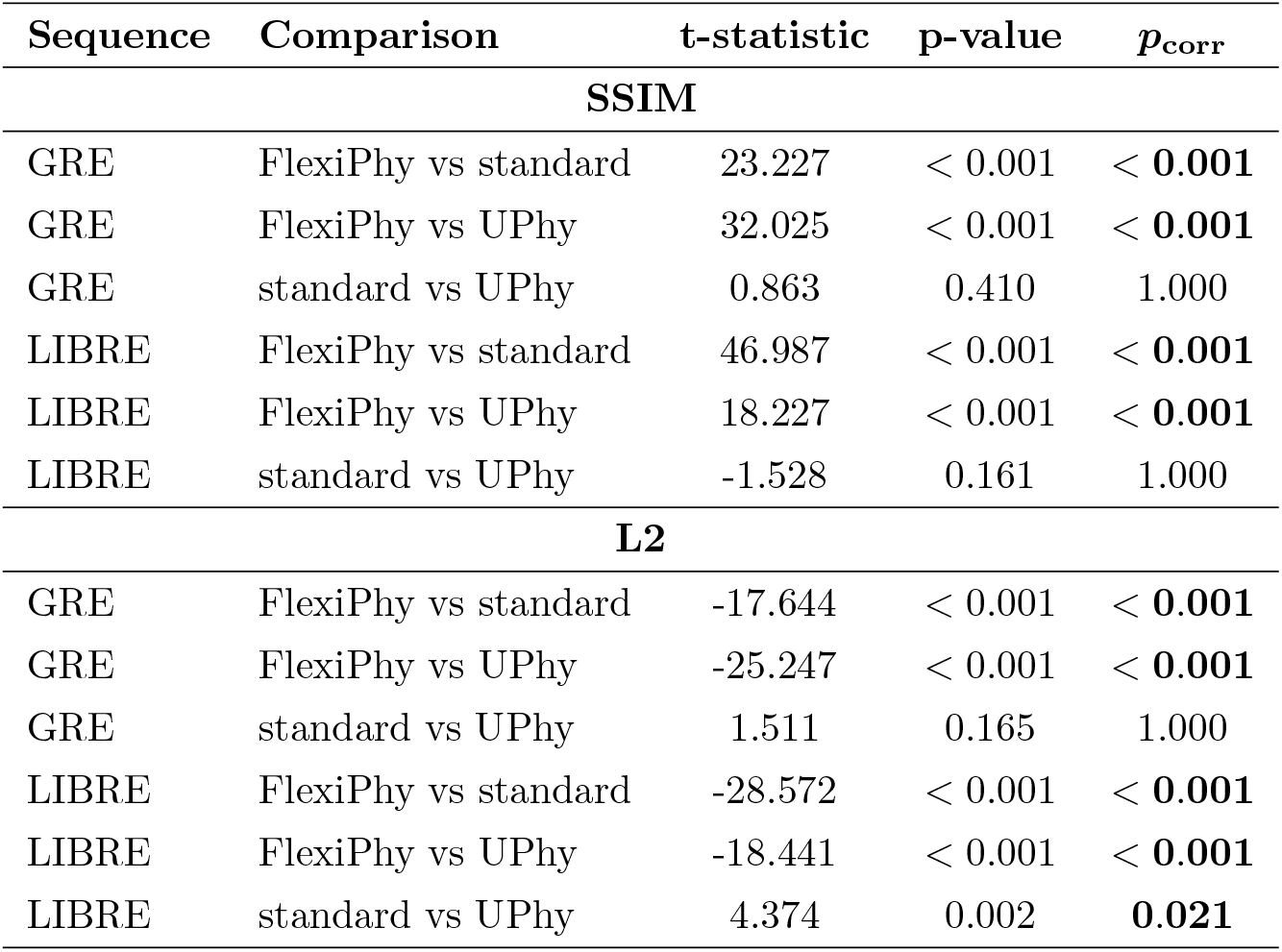
Statistical comparison of image-quality metrics across acquisition trajectories for GRE and LIBRE GRE sequences. Paired t-tests were performed on subject-level scores obtained by averaging the four sequential bins metric values for each subject, trajectory, sequence, and metric. All pairwise trajectory comparisons were tested separately for SSIM and relative L2 distance. P-values were corrected for multiple comparisons using the Bonferroni method across the 12 planned tests (3 trajectory comparisons × 2 metrics × 2 sequences), with *p*_corr_ = min(12*p*, 1). Corrected p-values satisfying *p*_corr_ < 0.05 are shown in bold. P-values smaller than 0.001 are reported as < 0.001. For relative L2 distance, lower values indicate improvement; for SSIM, higher values indicate improvement.

### 3.2 | Dynamic images: eyes

Figure 7 presents a cropped view of the eye region for the dynamic eye imaging reconstructions of the LIBRE GRE data across the four temporal bins. The standard phyllotaxis and UPhy trajectories produced structured ringing artifacts that propagated directly into the orbital region and varied across temporal bins. These artifacts partially obscured anatomical details and introduced temporal inconsistencies.

The FlexiPhy trajectory still presents some motion artifacts that are affecting the image quality, but substantially reduces ringing artifacts, leading to more homogeneous temporal reconstructions and improved visualization of the eye and surrounding orbital structures.

## 4 | DISCUSSION

In this work, two limitations of the classical 3D radial spiral phyllotaxis trajectory were investigated: the non-uniform SDSD and the emergence of ringing artifacts during sequential binning. To address these limitations, two new trajectory designs were introduced, namely UPhy and FlexiPhy, and their performance was evaluated in vivo using T1-weighted GRE and LIBRE acquisitions.

UPhy addresses the first limitation by modifying the polar-angle formulation of the standard spiral phyllotaxis trajectory to improve the density distribution of readout directions. This change is relevant for most applications, since non-uniform angular sampling can lead to direction-dependent variations in effective field of view, spatial resolution, and sampling efficiency ^32,34,33,28,29,30,31^. In this sense, UPhy provides an improved sampling pattern on the sphere while preserving the key properties of the standard spiral phyllotaxis trajectory.

However, visual inspection of the in vivo results indicates that improving the trajectory SDSD alone is not sufficient to remove artifacts associated with sequential binning. In sequentially binned reconstructions, image quality is determined not only by the global distribution of all acquired readouts, but also by the distribution of the subset assigned to each temporal bin. UPhy, as the standard phyllotaxis, preserves the coupling between polar and azimuthal ordering. This coupling induces a structured clustering of samples when consecutive interleaves are grouped into temporal bins, resulting in concentric k-space gaps, leading to r. This likely explains why UPhy improves theoretical angular uniformity without producing a corresponding reduction of the ringing artifacts observed in the binned reconstructions. This interpretation is supported by the statistical analysis in Table 2, where comparisons between UPhy and the standard phyllotaxis trajectory were generally not significant, with the sole exception of the LIBRE relative L2 metric.

These observations motivate a trajectory design that preserves the improved global sampling properties of UPhy while specifically addressing the non-uniform coverage of temporally contiguous subsets. FlexiPhy was designed for this purpose. By decoupling the polar-angle ordering from the azimuthal golden-angle progression through a randomized permutation of interleaves polar angles, FlexiPhy reduces systematic clustering within sequential bins while maintaining the local gradient behavior and interleaving structure of the standard trajectory. The improved image quality observed with FlexiPhy, also reflected in the statistical tests results (Table 2), therefore supports the interpretation that the dominant source of degradation in sequential temporal-bin reconstructions is the organization of readout directions across acquisition time.

Importantly, the randomized construction of FlexiPhy provides high statistical confidence that readout origins remain well distributed and free of large gaps across binning conditions. In this sense, FlexiPhy extends the flexibility of the standard phyllotaxis trajectory to arbitrary retrospective binning strategies. Because FlexiPhy incorporates the polar-angle definition introduced in UPhy, it improves the global distribution of readout directions while preserving small angular differences between successive readouts, limiting rapid gradient changes that could otherwise increase eddy-current artifacts, and retaining golden-angle spacing between successive interleaves.

These findings have practical implications for radial MRI applications that reconstruct images from consecutive or partially overlapping subsets of readouts. In applications such as dynamic imaging, motion-resolved reconstruction, rigid and non-rigid motion correction ^7^, functional MRI ^16^, and eye imaging ^12,13,14^, temporal subsets are often reconstructed independently or used to estimate motion parameters ^15^. In these settings, FlexiPhy is preferable to both the standard phyllotaxis trajectory and UPhy, because it directly targets the temporal-subset sampling problem that affects reconstruction stability.

The subject-level observations also highlight that differences in scan orientation can change the spatial manifestation and severity of the ringing pattern. The observation of similar artifact behavior at an independent imaging site using a different sequence development framework further supports the relevance of the proposed modification ^46^.

Several limitations should nevertheless be acknowledged. First, the present study considered only image-quality metrics assessing visual artifact reduction. Future work should investigate the impact of FlexiPhy on downstream tasks such as motion-estimation accuracy, functional sensitivity, or clinical diagnostic performance. Second, while this work specifically focused on sequential binning, many radial MRI applications rely on pseudo-random physiological binning strategies ^2,3,4,5,6,8,9,10,11^, such as respiratory- or cardiac-resolved imaging. In these contexts, the relative benefits of UPhy and FlexiPhy may differ and deserve dedicated investigation.

Finally, the proposed trajectory designs could be combined with recently introduced pole-to-pole spiral phyllotaxis trajectories ^35^, which acquire readouts from both hemispheres of k-space to mitigate artifacts arising from phase inconsistencies and gradient-system imperfections. Investigating whether the improvements in sampling uniformity and binning robustness provided by UPhy and FlexiPhy can be synergistically combined with the phase-error compensation properties of pole-to-pole acquisitions represents an interesting direction for future work. Such a combination could also extend the applicability of FlexiPhy to half-spoke acquisition techniques, including UTE and ZTE imaging.

## 5 | CONCLUSIONS

In this study, two modifications of the classical 3D radial spiral phyllotaxis trajectory were introduced. UPhy, a reformulation of the polar-angle definition, yielding a uniform SDSD. FlexiPhy is a trajectory redesign improving robustness to sequential binning by decoupling polar and azimuthal ordering while preserving the golden angle interleaving structure. Experiments performed in vivo using T1-weighted GRE and LIBRE acquisitions demonstrated that FlexiPhy substantially reduces ringing artifacts in sequential temporal reconstructions and significantly improves image similarity metrics relative to both the standard phyllotaxis and UPhy trajectories. Overall, FlexiPhy preserves the spiral phyllotaxis practical advantages while providing a more uniform SDSD and substantially improving robustness for dynamic and retrospectively binned MRI applications. FlexiPhy therefore represents a flexible and efficient alternative to the standard spiral phyllotaxis trajectory.

## 6 | ACKNOWLEDGEMENTS

We thank all contributors and organizations whose support has made this project possible. All scientific content and interpretations are the authors’ own. This work was supported by the Swiss National Science Foundation (SNSF): M.L., Y.J., J.B. were supported under grants #220433, #229214 to Prof. Franceschiello. J.B. was also supported under grant #129816IA-RECHERCHE23-19 from HES-SO University of Applied Sciences and Arts Western.

J.D. was supported under grant 200021 215336 to Prof. Delabays.

The Sense Innovation and Research Center (Grant: KiCk fMRI, PI: Prof. Franceschiello & Prof. Dr. Juliane Schneider).

OpenAI’s LLMs were used during the preparation of this manuscript for language editing and proofreading, and in limited cases for code-related assistance, primarily for minor refactoring, generation of code for plotting, and documentation generation.

We acknowledge the resources and expertise provided by the CIBM Center for Biomedical Imaging.

## 7 | CODE & DATA AVAILABILITY

Together with the paper we provide several resources:

1. Standard Operating Procedures for data acquisition and processing: https://mattechlab.github.io/sops/flexyphy/data-collection/data-collection_flexyphy_protocol/
2. Interactive viewer for the reconstructions, to visualize the images for all the subjects: https://mauroleidi.github.io/flexiphy_html_viz/viewer/index.html
3. The code repository: https://github.com/MattechLab/FlexiPhy
4. The Data: We are currently working to make the dataset publicly available. However, its large size (over 800 GB) has made this process more challenging than anticipated. In the meantime, we have proceeded with the manuscript submission for peer review and plan to include a link to the public data repository before any eventual publication.

