## Supplementary Materials for "Improved 3D Radial Phyllotaxis Trajectories for Uniform Density Distribution of Readout Directions and Sequential Binning"

### 8 | SUPPLEMENTARY MATERIAL

#### 8.1 | Construction of UPhy: Proof of uniformity

A spherical cap is a subset of a sphere whose boundary is defined by a circle. The goal of this section is to characterize the spatial density of readouts generated by the standard phyllotaxis trajectory origins and compare it with the proposed UPhy formulation. To this end, we compute the density of readouts origins within a spherical cap as a function of the polar angle and show that the standard phyllotaxis trajectory produces a non-uniform distribution, whereas UPhy yields a constant density over the hemisphere. Figure 9 shows the spherical cap $(0, \theta)$. Its area is $A_\theta$, and the amount of points it contains as $P_\theta$. If $N$ readouts are placed on the surface of the hemisphere according to the standard phyllotaxis spiral trajectory, then let us compute the density $\Delta(\theta) = \frac{P_\theta}{A_\theta}$ of points in the spherical cap in function of $\theta$.

$$P_\theta = \sum_{n=1}^{N} \mathbf{1}(\theta > \theta_n)$$

where $\mathbf{1}(\theta > \theta_n)$ is an indicator variable that has value 1 if $\theta > \theta_n = \frac{\pi}{2}\sqrt{\frac{n}{N}}$ and 0 otherwise, hence indicates if readout $y_n$ is in $P_\theta$. The amount of points sampled in the cap is then:

$$\sum_{n=1}^{N} \mathbf{1}(\theta_n < \theta) = \sum_{n=1}^{N\left(\frac{2\theta}{\pi}\right)^2} 1 = N\left(\frac{2\theta}{\pi}\right)^2 = 4N\left(\frac{\theta}{\pi}\right)^2 = P_\theta.$$

*Remark 1.* Strictly speaking, the expression should be written as

$$\sum_{n=1}^{N\left(\frac{2\theta}{\pi}\right)^2} 1 = \lfloor N\left(\frac{2\theta}{\pi}\right)^2 \rfloor.$$

However, since the choice of $\theta$ is arbitrary, it may be selected such that the right-hand side is an integer. Even when this is not the case, the resulting discrepancy remains negligible for the present purposes, as illustrated in Figure 10 .

On the other hand, the area of a spherical cap is:

$$A_\theta = 2\pi R h_\theta$$

where $h_\theta = R(1 - \cos(\theta))$. Thus,

$$A_\theta = 2\pi R^2(1 - \cos(\theta)).$$

The density $\Delta(\theta)$ can now be computed. It is shown to be an increasing function of $\theta$, in particular, not constant on the surface of the sphere.

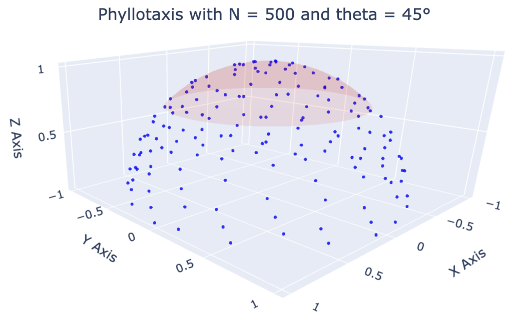


**FIGURE 9** In red you can observe a spherical cap of area $A_\theta$ with $\theta = 45°$. The goal is that the density $P_\theta / A_\theta$ of points in a given spherical cap is independent on $\theta$.

$$\Delta(\theta) = \frac{P_\theta}{A_\theta} = \frac{4N\theta^2/\pi^2}{2\pi R^2(1 - \cos(\theta))} \quad (1)$$

$$= \underbrace{\frac{2N}{\pi^3 R^2}}_{\text{constant}} \frac{\theta^2}{1 - \cos(\theta)} = \kappa \cdot \frac{\theta^2}{1 - \cos(\theta)} \quad (2)$$

which is strictly increasing on $(0, \frac{\pi}{2}) \ni \theta$ as shown in Figure 10 . In other words, this means that the sampling strategy places more readouts near the equator. The values range from $\Delta(\theta = 0) = 2\kappa$ to $\Delta(\theta = \frac{\pi}{2}) \approx 2.467\kappa$.

On the other hand, since the area of the cap equals: $A_\theta = 2\pi R^2(1 - \cos(\theta))$, the choice of an alternative polar angle $\tilde{\theta}_n = \arccos(1 - \frac{n}{N})$ yields the following amount of points:

$$\sum_{n=1}^{N} \mathbf{1}(\tilde{\theta}_n < \theta) = \sum_{n=1}^{N(1-cos(\theta))} 1 = N(1 - cos(\theta)) = \tilde{P}_\theta.$$

and therefore the density:

$$\frac{\tilde{P}_\theta}{A_\theta} = \frac{N(1 - \cos(\theta))}{2\pi R^2(1 - \cos(\theta))} = \frac{N}{2\pi R^2}$$

which is indeed constant.

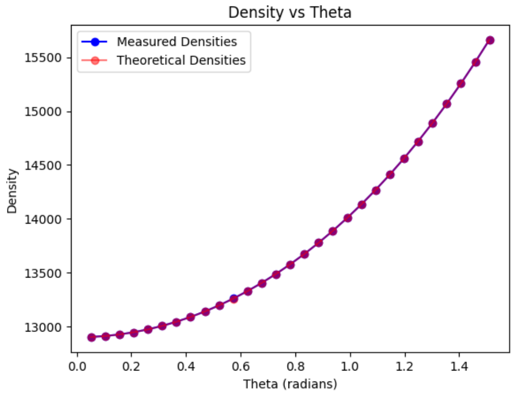


**FIGURE 10** Comparison between the theoretical and empirical densities of readouts inside the spherical cap $(0, \theta)$ for the standard phyllotaxis sampling on a hemisphere. The empirical density is obtained numerically by generating the sampling pattern using the standard phyllotaxis formulation, and then counting the sampled points contained in the cap and dividing by its area, while the theoretical density is given by $\Delta(\theta) = \kappa \frac{\theta^2}{1-\cos(\theta)}$. The agreement between both curves validates the analytical computation and shows that the density increases with $\theta$, indicating a higher concentration of readouts near the equator.

#### 8.2 | Multicentric, Multi-Sequence Development Tool and in Phantom Validation

This subsection presents preliminary results acquired for our 2025 ISMRM abstract[46]. These data were obtained using a sequence implemented in IDEA and were acquired at a different site from the study data presented in this work. The dataset includes a Siemens water bottle phantom and 3 in vivo acquisitions. Notably, the observed artifacts closely resemble those seen in Subject 1 of the study dataset, suggesting that they are associated with a different acquisition orientation.

#### 8.3 | Comparison with the Golden Step Trajectory

A previously proposed solution to mitigate ringing artifacts is the Golden Step trajectory[28]. For certain combinations of trajectory parameters, this approach can provide a good distribution of sequential interleaves origins, as illustrated in the top panel of Figure 11 . However, the method does not preserve the golden-angle property of the standard phyllotaxis design. Consequently, consecutive interleaves are no longer separated by the golden angle, as shown in Figure 12 .

More importantly, the performance of the Golden Step trajectory is highly dependent on the choice of trajectory parameters. For example, in the case where $F$ is a Fibonacci number, consecutive interleaves cluster together rather than being evenly distributed across the sphere. This behavior creates large gaps in groups of interleaves and compromises the flexibility of the trajectory, as illustrated in Figure 11 and Figure 12 .

In contrast, Flexiphy preserves the golden-angle property while avoiding the clustering observed with the Golden Step approach, resulting in improved sampling properties across trajectory parameters.

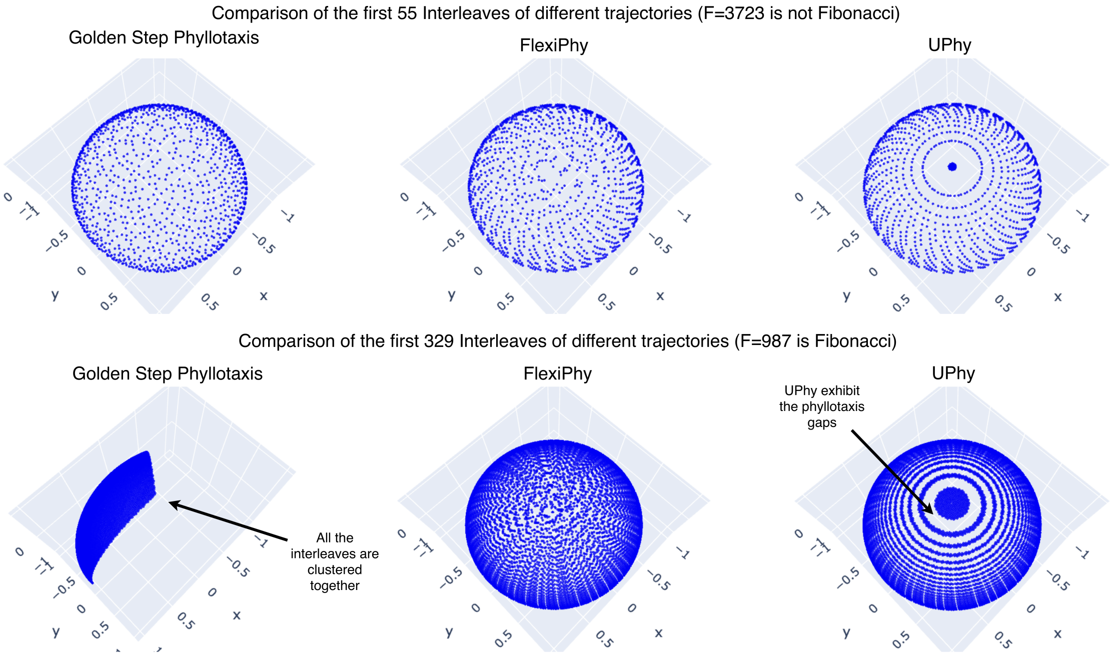


**FIGURE 11** On the top panel we observe the first 55 interleaves and $F$ is not a Fibonacci number. We can observe that the Golden Step approach distributes points quite well for this trajectory parameter combination. On the bottom panel, we observe the first 329 interleaves and $F$ is chosen to be a large Fibonacci number. We observe that for $F$ large Fibonacci number, the Golden Step approach clusters interleaves together. Gaps are visible in the uniform phyllotaxis design. The amount of points in each interleaves $P = 22$ for both panels.

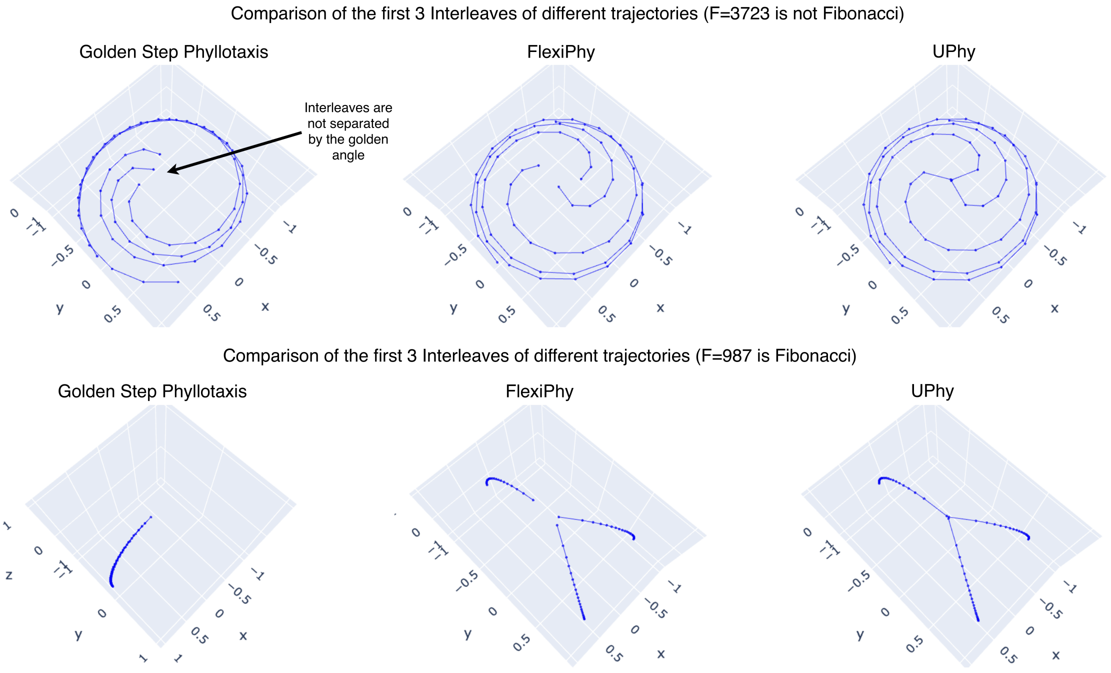


**FIGURE 12** Comparison of the first 3 interleaves of the Golden Step Phyllotaxis with the standard phyllotaxis and Flexiphy. On the top panel $F$ is not a Fibonacci number. We can observe that the Golden Step approach does not maintain the golden angle property and therefore interleaves are not separated by the golden angle. On the bottom panel, $F$ is chosen to be a large Fibonacci number. We observe that for $F$ large Fibonacci number, the Golden Step approach clusters interleaves together. Gaps are visible in the uniform phyllotaxis design. The amount of points in each interleaves $P = 22$ for both panels.

##### 8.4 | Metrics used in section 3 taken from the work of Piccini & al.

In Figure 5 , two features of trajectories were presented to illustrate how UPhy improves the uniformity of the phyllotaxis spiral in non-radial directions. Both metrics were defined and used in[1]. First, the average distance between two successive points of an interleaf is simply defined as

$$m_{dist} = \frac{1}{P-1}\sum_{j=1}^{P-1} d(y_j, y_{j+1}).$$

Remark that by symmetry, this mean distance should hardly depend on the choice of the interleaf, hence our decision to only compute this indicator for the first one.

The second one represents the uniformity of points on the surface of the half-sphere, by calculating the relative standard deviation of the distance between a point $i$ and its four closest neighbours. The metric consists in the average of these RSDs over all points $i$ of the trajectory, that is,

$$\text{RSD} = \frac{\sqrt{\frac{1}{4N}\sum_{i=1}^{N}\sum_{j=1}^{4}(d_{i,j}-\mu_d)^2}}{\mu_d}\cdot 100$$

where $\mu_d$ is the global mean distance from every point to its four closest neighbours:

$$\mu_d = \frac{1}{4N}\sum_{i=1}^{N}\sum_{j=1}^{4} d_{i,j}.$$

UPhy and the standard phyllotaxis trajectory were evaluated with these two metrics, and summarized the results in Figure 5 .

##### 8.5 | Random permutation

###### 8.5.1 | Why choosing $\sigma$ at random?

In section 2.3, the polar and azimuthal angles were decoupled via a permutation $\sigma$ of the numbers $\{1, 2, \ldots, F\}$. Indeed, in the standard model, these angles are coherent, meaning that the interleaf that would correspond to the $i$-th interleaf of a planar phyllotaxis spiral in the $xy$-plane is the $i$-th highest with respect to the $z$-axis. Our decoupling breaks this coherence, so that now the $i$-th interleaf in the $xy$-plane is the $\sigma(i)$-th highest.

The choice of $\sigma$ at random among the $F!$ existing permutations seems an uncertain decision. However, to our knowledge, every attempt at taking a structured $\sigma$ (such as[28]) leads to undesirable patterns, see Figure 12 . This study failed many attempts at finding a deterministic way to select a decoupling that was robust across trajectory parameters (avoiding patterns and gaps).

The number of existing permutations $F!$ is generally huge: a typical value in practice is $F = 2440$, yielding $F! \approx 4.4 \cdot 10^{7'207}$. As a consequence, the probability that $\sigma$ is not suitable for image reconstruction is negligible. For example, consider a given bin $B$ containing 20% of the data point, that is, 488 of the 2440 interleaves. If the readouts were placed according to the FlexiPhy pattern, then the probability for $B$ to contain no interleaf in the lowest 10% of all interleaves would be

$$(2440-488)!\frac{(2440\cdot 0.9)!}{(2440\cdot 0.9-488)!}/2440! \approx 8.45\cdot 10^{-26}. \quad (3)$$

It is hereby statistically safe to assume that the interleaf $B$ spans a representative variety of the interleaves.

*Remark 2.* Formula (3) is obtained as follows. First, the total number of permutations is $F!$. Moreover, the permutations $\sigma$ that would lead $B$ to miss on 10% of the interleaves are the elements of the set $S_{miss} := \{\sigma \in S_F \mid \sigma(i) \leq M := 0.9 \cdot F$ for any $i \leq |B|\}$, the cardinality of which is $\frac{M!}{(M-|B|)!}(F-|B|)!$. Indeed, the elements of $S_{miss}$ are elements of $S_F$ that match the first $|B|$ elements to any of the $M$ spots, and the last $(F-|B|)$ elements to the remaining spots. The fraction of favorable cases then equals $\frac{|S_{miss}|}{|S_F|}$, which indeed is close to $8.45\cdot 10^{-26}$ when $F = 2440$ and $|B| = 0.2 \cdot F = 488$.

###### 8.5.2 | Precautions of use linked to random permutation

The permutation used during acquisition must also be available during reconstruction in order to recover the correct trajectory. This requirement can be problematic when the permutation is generated in an old software environment, for example using C++ versions without standardized random-number generation. In our case, this was one of the main reasons for implementing the sequence trajectory directly in Pulseq.

**How to cite this article:** Leidi M., Délitroz J., Peper E., Jia Y., Barranco J., Ledoux J.-B., Romanin L., Bastiaansen J., Schneider J., and Franceschiello B. (2026), Improved 3D Radial Phyllotaxis Trajectories for Uniform Density Distribution of Readout Directions and Sequential Binning, *J. Magn. Reson. Imaging, 2026;00:000.*
